# Transfer learning applied in predicting small molecule bioactivity

**DOI:** 10.1101/2025.01.09.632140

**Authors:** Li Tian, Hugh Hongkai Huang, Eric Martin, Judy Ying Wei, Shaoyong Xu, Junzhou Huang

**Affiliations:** Laboratory for Synthetic Chemistry and Chemical Biology Limited (LSCCB), Hong Kong SAR; Tencent AI Lab, Shenzhen, P.R.China; Novartis Institute for Biomedical Research, California, United States; College of Computer Science and Technology, Zhejiang University, P.R.China; Computer Science and Engineering department at the University of Texas at Arlington, United States

## Abstract

Despite over a half-century of effort by computational chemists, developing accurate empirical QSAR (Quantitative Structure-Activity Relationships) models for predicting bioactivity directly from chemical structure has remained elusive. The difficulties have been especially pronounced for virtual screening, finding new active compounds substantially different from the known chemical matter used to train the models. Recent breakthroughs have been achieved by employing transfer of learning across huge numbers of bioactivity assays, greatly increasing the amount and diversity of chemical and biological information that informs each model. An early example was Profile-QSAR (pQSAR), a 2-level stacked model, where level-2 PLS (Partial Least Squares) models characterize compounds by their profile of bioactivity predictions from individual level-1 random forest regression QSAR models built on up to 10,000+ other assays. This study introduces metaNN, a meta-learner that trains deep neural networks (DNN) for each individual assay initialised from a well-generalized consensus DNN optimized across all assays. Comparison of the results suggested that while Profile-QSAR and metaNN perform similarly overall, metaNN works slightly better for smaller assays which were well-predicted by the consensus DNN; whereas pQSAR struggled more with smaller assays, due to the large number of level-1 models but was less sensitive to similarity to an overall consensus. An ensemble average of both methods combined the strengths of each, working better than either alone. The similar performance of the 2 largely orthogonal algorithms raises questions about whether we are approaching a limit of prediction accuracy in transfer learning, for this application scenario.

## Introduction

Recently, significant interest has been focused on applying artificial intelligence (AI) algorithms to predict small molecule bioactivity and protein-binding affinity. Structure-based methods are supposed to learn the physics of interactions between the small molecule ligands and the protein pockets, and thus provide accurate prediction with no training data. However, for many drug discovery projects, the target protein or even biological mechanism is not completely clear, or a high-resolution structure of the binding complex may not be available. These factors limit the applicability of structure-based prediction methods. Furthermore, physics-based methods typically ignore many important factors that influence activity: concentrations of cofactors, post-translational modifications, allosteric modulations such as a protein complex formation, identity of second substrates (e.g., the particular peptide substrate for a protein kinases ATP-site inhibitor), and all the complexity of cellular assays. These factors complicate the states of the protein and limit the accuracy or sometimes even the relevance of the structure-based prediction. One alternative approach starts from the compounds’ chemical structures and bioactivity data, i.e., the “ligand-based” data sets. These data sets are much more widely available, including functional and cellular assays. These data sets can be used to build a ligand-based model for activity prediction without prior knowledge of the target protein or even the biological mechanism. How to best leverage this significantly bigger data pool and convert these data sets to knowledge and predictive models, is of great interest and has wide application in drug discovery.

Chemical informaticians have been building ligand-based models for decades, from very traditional QSAR models based on physicochemical properties to random forest models based on molecular fingerprints. Some of these models may achieve impressive prediction accuracy on random test sets, yet fail to demonstrate comparable accuracy in realistic applications such as virtual screening to identify novel hit compounds or off-target prediction. Although the reasons behind this limitation may be complicated, the “stretched” extrapolation of the “local model” trained from small data sets to much larger chemical space for prediction, may account for the major contribution. On the other hand, the limitations rooted from sparse known data have also been well studied by machine learning (ML) scientists: how to effectively learn from a very small amount of data and apply to larger prediction sets. Recently, several attempts have been made leveraging knowledge transfer to remedy the lack of bioactivity data, and thus more effectively train predictive models. These studies demonstrate two major strands of algorithms: transfer learning (*1*) and multitask learning(*2*). Transfer learning aims to improve the bioactivity prediction accuracy of a target assay via transferring the knowledge learned from previous source assays; whereas multitask learning jointly learns all the assays together, without differentiating source and target assays. Girschick et al. (*3*) proposed to learn a distance measure that reflects the similarities between instances and transfer that measure to the target assy. Profile-QSAR (pQSAR) [4] uses a 2-level stacked model (*4*), where a compound is characterized by its profile of bioactivity predictions from QSAR models built on many other assays. Similar to pQSAR, Alchemite also uses stacked models, but is based on neural networks rather than older random forest and PLS algorithms.(*5*) Multi-target QSAR models were built on a subset of the human kinome with taxonomy-based multitask methods proposed in a recent study(*6*). Recent advances in deep neural networks have motivated deep transfer learning (DTL) and deep multitask learning (DMTL) studies, where deep neural networks with greater expressiveness are taken as predictive models. The most prevalent DTL strategy is fine-tuning--a neural network that predicts an activity or property is pre-trained on all source assays and fine-tuned to the target assay (*7, 8*). Another state-of-the-art method(*9*) learns an attention function that measures the similarity between instances and transfers the function to the target assay, which shares the spirit of Girschic’s work (*3*). Despite its superiority over single-assay neural networks, DTL ignores the difference between different assays. Instead, DMTL learns different predictors for different assays, while sharing the feature extractor at low-level layers among all assays. By virtue of the high predictive capacity of deep neural networks, and knowledge sharing between assays, both Dahl et al.(*10*) and Ramsundar et al.(*11*) have reported that DMTL outperforms single-assay neural networks. To further improve the effectiveness of DMTL with fine-grained knowledge sharing, binding domain sequence similarity(*12*) and amino-acid sequence similarity(*13*) have been leveraged to model the relationship between targets. The downside of DMTL lies in the risk of an assay-specific predictor being overfitted to extremely scarce bioactivity data.

In this paper, we seek an approach to address the problems of both DTL and DMTL, where a neural network is globally shared across all assays, while it is explicitly trained to adapt to each in only a few steps with low risk of overfitting. Transferring and adapting the global neural network quickly trains the model for any specific target assay. Here we describe a new meta-learning algorithm, named metaNN, discuss the similarity and dissimilarity in predictions between metaNN and pQSAR, analyze the results in detail, and show that a combination or ensemble method achieves even higher accuracy.

## 2. MetaNN algorithm

Our proposed algorithm, metaNN, is inspired by the recent success of gradient-based meta-learning(*14*), which leverages the meta-knowledge learned from previous tasks to facilitate the learning of a novel task. This paradigm creates task-specific neural networks, while requiring minimum learning for each individual task, thus minimizing the risk of overfitting given limited training data. Figure 1 presents an overview of the workflow of gradient-based meta-learning algorithms for the problem of bioactivity prediction: 1) we train a neural network whose initialization for the weights is shared across all *n* source assays; 2) for a source assay, e.g. source assay 1, we build the initialized neural network for its training-set compounds on bioactivity values for its assay; 3) we evaluate the adapted neural network on test-set compounds of each source assay, so that the error rates from all source assays are averaged and back-propagated to improve the generalization capability of the initialization; 4) the learned initialization, called meta-initialization, is transferred to facilitate the training of the target assay.

**Figure 1.**
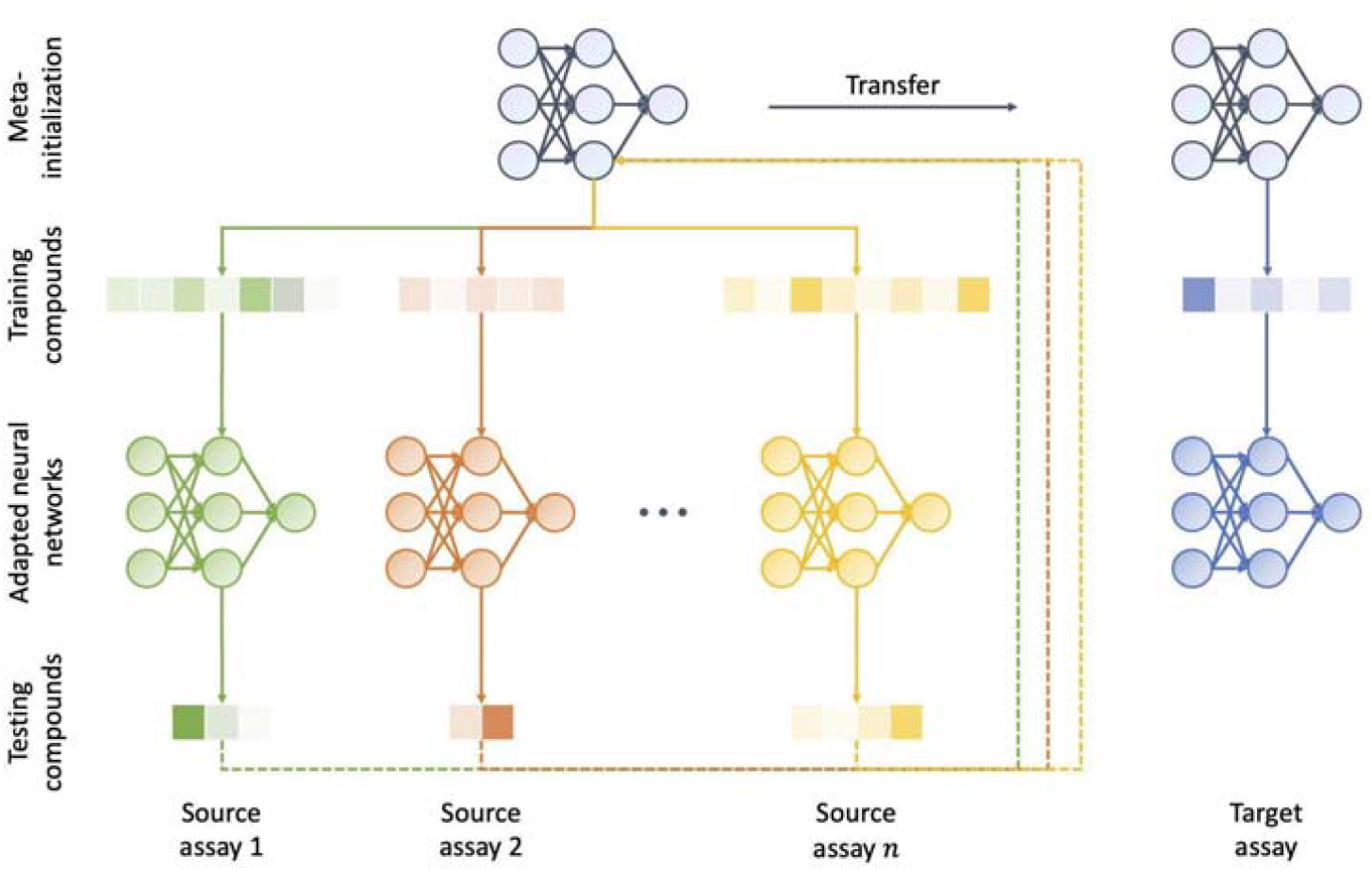
Pictorial illustration of the workflow of the metaNN algorithm.

Mathematically, we denote each assay as *T*_*i*_ with its training set as 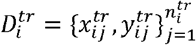 and testing set as 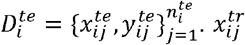 denotes the *j*-th compound for the *i*-th assay, and 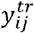 is its pIC_50_ activity value. 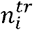 and 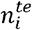 are the numbers of training and testing compounds in the *i*-th assay, respectively. Suppose that the neural network is represented by *f*_*W*_, where *W* denotes all weights for the neural network. We evaluate the performance of the neural network with the squared loss between the predicted and experimental bioactivity values, i.e., 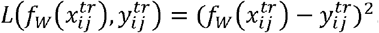. *W*_0_ is the meta-initialization for the neural network *f*_*W*_ which we aim to learn. The workflow of gradient-based meta-learning, therefore, boils down to the following two alternating optimization problems.

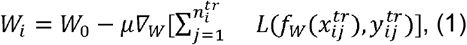

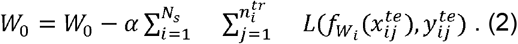

The first equation corresponds to step 2) where all training compounds are used to adapt the meta-initialization *W*_0_ to be assay-specific via several steps of gradient descent, while the second equation for step 3) learns a well-generalized meta-initialization such that the error rates of adapted neural networks starting from *W*_0_ on test compounds of all source assays are minimized. *µ* and *α* are the learning rates for updating *W*_*i*_ and *W*_0_, respectively. Provided with a target assay *T*_*t*_, the weights for its specific neural network are learned in a manner similar to the first equation, i.e.,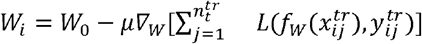. Considering the high generalization capacity of the optimized meta-initialization *W*_0_ for source assays, fast learning and satisfactory predictive accuracy of *W*_*t*_ on the target assay can be anticipated.

In the first set of calculations, this pipeline of gradient-based meta-learning did not achieve the accuracies we expected on the target assay. This implies that the learned meta-initialization does not generalize well. A well-qualified meta-initialization *W*_0_ should be consistently under-performing on either the training or the test set, leaving sufficient room for training compounds to learn the assay-specific weights *W*_*i*_ that suffice to achieve high prediction accuracies for both sets. Empirical results in Figure 2 show that even without adaptation to each individual assay, the meta-initialization *W*_0_ achieves extremely high accuracies on the test sets of all source assays, but quite low accuracies on the training sets. This suggests that the meta-initialization *W*_0_ simply memorizes the test sets of all source assays, rather than being a well-generalized starting point for the neural network to be optimized for each source assay through its training set.

**Figure 2.**
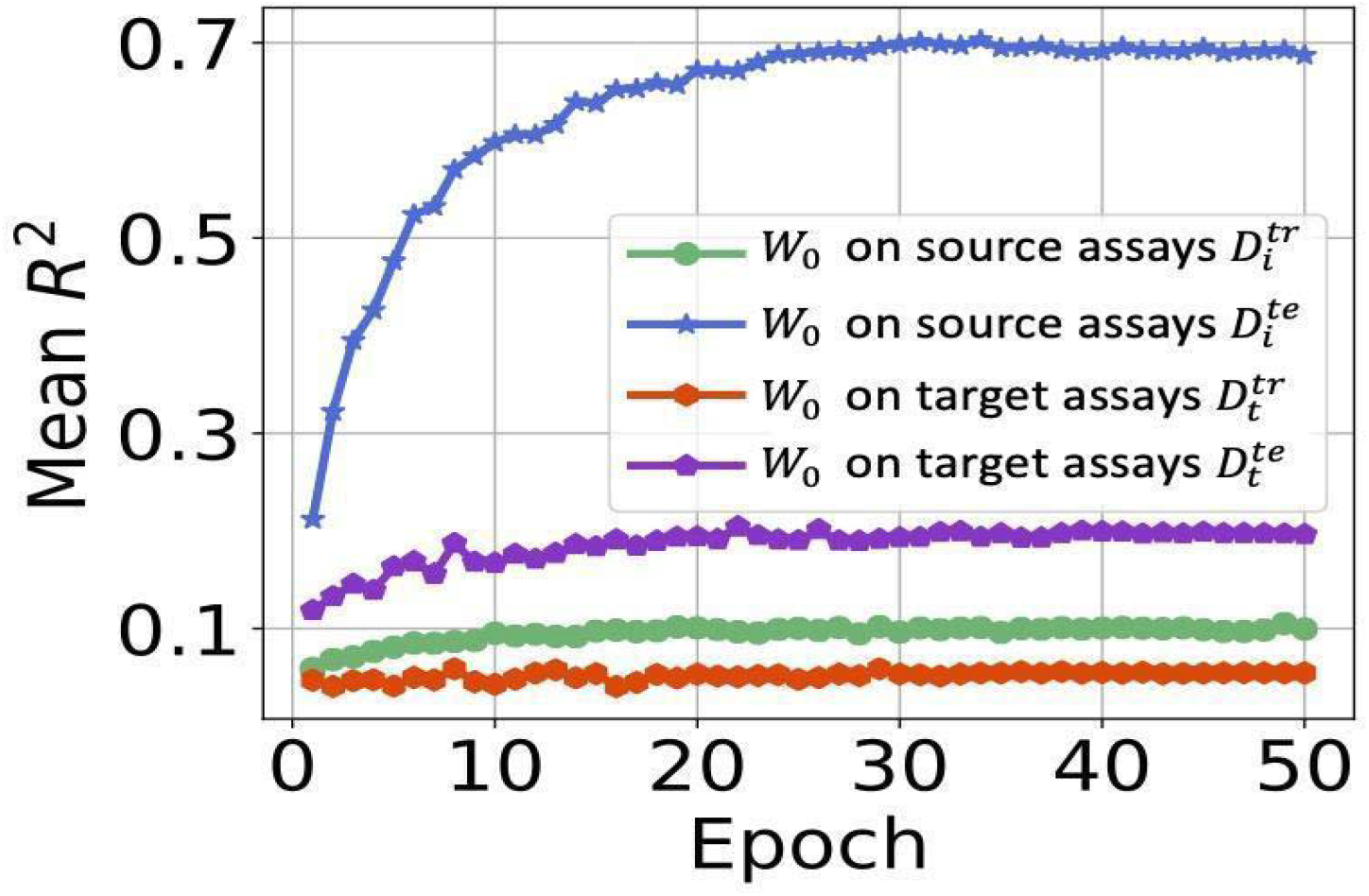
Inconsistent behaviors of *W*_0_ on training sets and testing sets.

We addressed this issue by producing more data to evaluate and update the meta-initialization in Eqn. (2). We list three desiderata for producing such a set of extra data: 1) the incorporation of such data would encourage consistent predictive performances of the meta-initialization across training and testing sets; 2) the data should contain more than the training set, as the training set has been used for training and fails to evaluate the generalization capability; 3) the data should contain more than the testing set, otherwise the memorization problem persists. Inspired by(*15*), we produce extra data by mixing both training and testing sets, where we mix-up not only hidden representations of compounds, but also their activity values. Specifically, for each randomly selected pair of training a compound 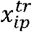 and a testing compound 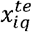 in the *i* -th source assay, we produce a new compound as

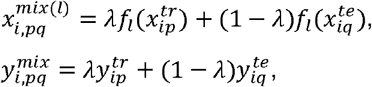

where *f*_*l*_ (·) is the *l*-th layer hidden representation by the neural network, and *λ* ∼*Beta*(*α, β*) is a randomly sampled coefficient from the Beta distribution. By sampling different pairs of training and testing compounds and sampling various values of *λ*, we produce a new collection of data that can be joined with the testing set to update the meta-initialization in Eqn. (2).

With the above-described algorithm, we used the same compound featurization as proposed by the pQSAR developers -- Morgan radius 2 substructure fingerprints of 1024 bits from RDkit v2018.03, trained the models, predicted bioactivities and evaluated prediction performance. Two data sets had been downloaded from ChEMBL and published in two earlier pQSAR papers: namely IC_50_s for 159 kinase assays(*16*) and for 4276 assays from all diverse families(*17*). The assay statistics of these two data sets are summarized in Table 1. The 159 kinase data set is smaller--focused on the most studied protein family in drug discovery. The 4276 assay set includes assays from 5 protein families, including kinases, proteases, GPCR, ion channels, nuclear hormone receptors, as well as phenotypic assays without clear targets registered. These two data sets both cover different assay formats, both biochemical and cellular, and the data maintains a wide dynamic range as a good reflection of real-life compound optimization projects.

**Table 1.** Summary statistics of the two datasets we evaluated.

| ChEMBL | assay cnt | total cmpd cnt | data cnt | data/cmpd | data/assay | protein families | target cnt |
| --- | --- | --- | --- | --- | --- | --- | --- |
| 159 kinase data set | 159 | 13k | 110k | 8.7 |  | kinase |  |
| 4276 all family data set | 4276 | 500k | 1.4m | 2.8 | 327 | 5 families + phenotypic |  |

In our calculations, we chose to use the same two public domain data sets for direct comparisons to pQSAR and easier access for other researchers in this field. When evaluating the model performance, we mainly use squared Pearson correlation coefficient between the experimental and predicted pIC_50_ values for the compounds in the external test sets, noted as R^2^_ext_. A detailed discussion on why squared Pearson correlation coefficient was preferred for virtual screening use-cases, and its comparison to other evaluation metrics such as RMSE and MAE, was presented in earlier publications(*18*). We also continued using the realistic training/test set splits discussed in the earlier papers(*16*), and used the exact published training test sets. When evaluating model quality across many assays, the median R^2^ _ext_, and the number of models with R^2^_ext_ >= 0.3, were used as the two most important indicators.

For metaNN calculation, the neural network for predicting pIC_50_ activity was a two-layer Multi-layer Perceptron (MLP) with 500 hidden neurons in each layer. Also, each fully connected layer was followed by a batch normalization layer and a non-linear activation of leakyReLU (negative slope is 0.01). We updated the meta-initialization in a batch-wise manner, with each batch consisting of 8 randomly selected source assays, i.e., *N*_*s*_ = 8. The learning rates to update the meta-initialization (i.e., *α*) and to learn assay-specific weights (i.e., *µ*) were 0.001 and 0.01, respectively. We set the Beta distribution to sample values of *λ* as *Beta*(0.5,0.5), i.e., *α* = *β* = 0.5. We iteratively ran 50 epochs to update the meta-initialization, with each epoch including 500 iterations, while we took only 5 gradient steps to quickly learn assay-specific weights for the neural network in Eqn. (1).

## 3. Results

As the pQSAR code was not yet published, we first programmed and reproduced the pQSAR 2.0 result, and then compared its predictions with our methods. We modeled 4266 of the 4276 assays, having removed 10 assays whose test set compounds all have the same experimentally measured values. Tables 2 and 3 summarize the results. For both the 159 and the 4266 data sets, metaNN shows very similar overall prediction performance as pQSAR 2.0, evaluated by various summarizing statistics of R^2^_ext_. For the 4266 data set, the median R^2^_ext_ reported in the original publication of pQSAR 2.0 for the 4266 assays was at 0.30. In reduced profile pQSAR, the authors only used the random forest models whose predicted pIC_50_s correlate (r^2^) with the experimental measurements in the training sets above select thresholds. Models were built using three r^2^ thresholds: 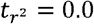 which corresponds to the full profile of all random forest models, 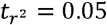 and 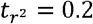. In our reproduced pQSAR 2.0 calculation, these 3 thresholds, 0.0, 0.05 and 0.2, provide R^2^_ext_ median at 0.26, 0.30, and 0.33, respectively. In max2 pQSAR(*17*), the authors used the test set R^2^_ext_ to select the best of the 3 models. The improvement brought by this model selection, (here named as max3) in Table 2, boosted the median R^2^_ext_ over the 0.2 threshold model from 0.33 to 0.41, and added 357 more successful models (R^2^_ext_>0.3). This reproduces the reported results from the publication. However, using the test set to choose among 3 models introduces a multiple-hypothesis testing issue, i.e. tripling the opportunities for chance correlations. The authors repeated the calculations with Y-scrambled data sets to demonstrate that even with the model selection strategy there were very few successful models simply achieved by chance. We likewise only observed an increase in the y-scrambled median R^2^_ext_ from 0.02 to 0.05. The number of y-scrambled models with R^2^_ext_>0.3 only increased by 107. We thus concluded that the published performance is very well reproduced and similar improvement over chance from model selection was observed.

**Table 2.**
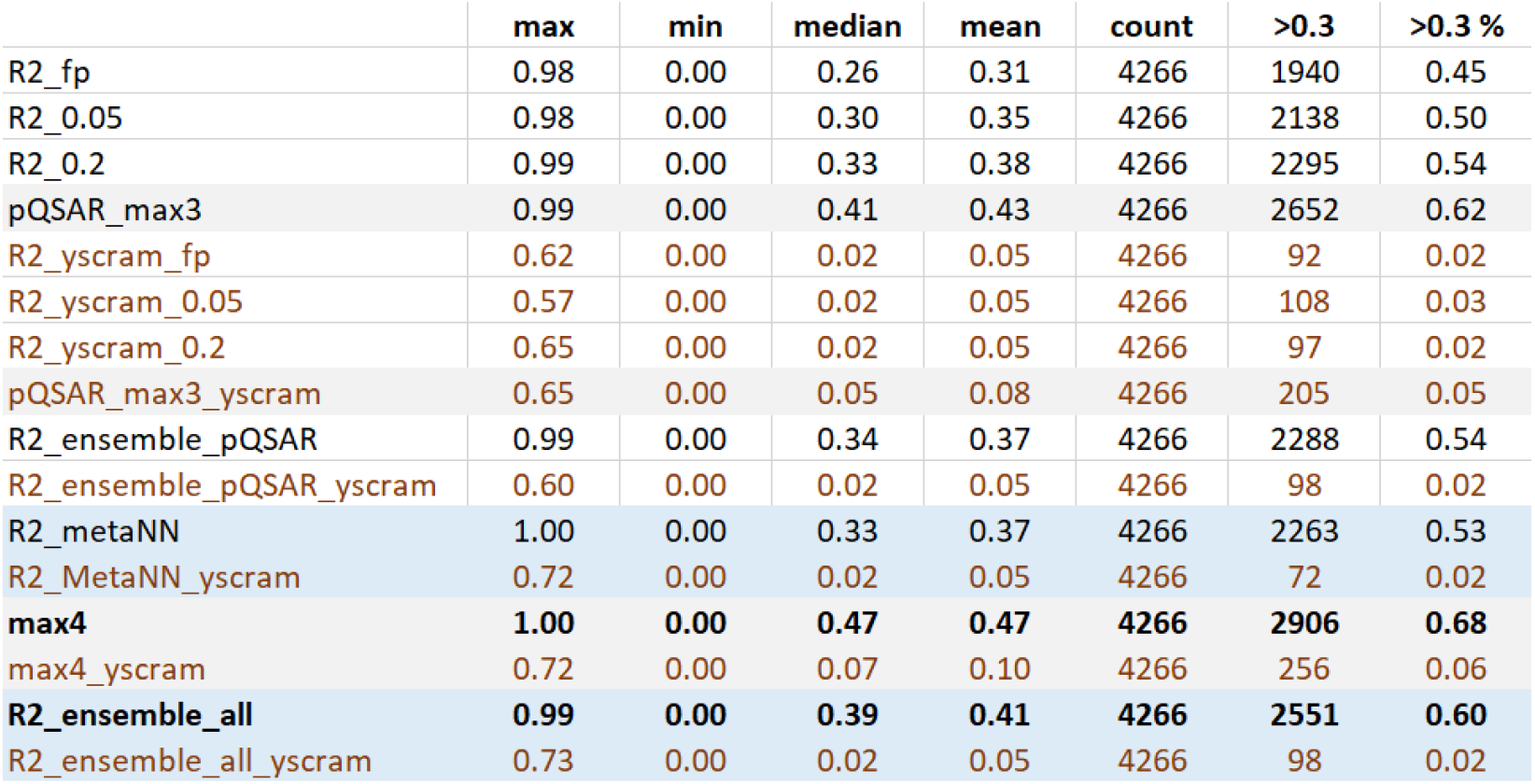
Summary of correlations R^2^_ext_ between experiment and prediction on 4266 ChEMBL assays for various multitask models.

|  | max | min | median | mean | count | >0.3 | >0.3 % |
| --- | --- | --- | --- | --- | --- | --- | --- |
| R2_fp | 0.98 | 0.00 | 0.26 | 0.31 | 4266 | 1940 | 0.45 |
| R2_0.05 | 0.98 | 0.00 | 0.30 | 0.35 | 4266 | 2138 | 0.50 |
| R2_0.2 | 0.99 | 0.00 | 0.33 | 0.38 | 4266 | 2295 | 0.54 |
| pQSAR_max3 | 0.99 | 0.00 | 0.41 | 0.43 | 4266 | 2652 | 0.62 |
| R2_yscram_fp | 0.62 | 0.00 | 0.02 | 0.05 | 4266 | 92 | 0.02 |
| R2_yscram_0.05 | 0.57 | 0.00 | 0.02 | 0.05 | 4266 | 108 | 0.03 |
| R2_yscram_0.2 | 0.65 | 0.00 | 0.02 | 0.05 | 4266 | 97 | 0.02 |
| pQSAR_max3_yscram | 0.65 | 0.00 | 0.05 | 0.08 | 4266 | 205 | 0.05 |
| R2_ensemble_pQSAR | 0.99 | 0.00 | 0.34 | 0.37 | 4266 | 2288 | 0.54 |
| R2_ensemble_pQSAR_yscram | 0.60 | 0.00 | 0.02 | 0.05 | 4266 | 98 | 0.02 |
| R2_metaNN | 1.00 | 0.00 | 0.33 | 0.37 | 4266 | 2263 | 0.53 |
| R2_MetaNN_yscram | 0.72 | 0.00 | 0.02 | 0.05 | 4266 | 72 | 0.02 |
| max4 | 1.00 | 0.00 | 0.47 | 0.47 | 4266 | 2906 | 0.68 |
| max4_yscram | 0.72 | 0.00 | 0.07 | 0.10 | 4266 | 256 | 0.06 |
| R2_ensemble_all | 0.99 | 0.00 | 0.39 | 0.41 | 4266 | 2551 | 0.60 |
| R2_ensemble_all_yscram | 0.73 | 0.00 | 0.02 | 0.05 | 4266 | 98 | 0.02 |

**Table 3.** Performance comparison for the 159 data set.

| R2ext | max | min | median | mean | count | >0.3 | >0.3 % |
| --- | --- | --- | --- | --- | --- | --- | --- |
| pQSAR_fp | 0.86 | 0.00 | 0.47 | 0.46 | 159 | 130 | 0.82 |
| pQSAR_0.05 | 0.90 | 0.00 | 0.47 | 0.45 | 159 | 126 | 0.79 |
| pQSAR_0.2 | 0.87 | 0.00 | 0.46 | 0.45 | 159 | 128 | 0.81 |
| pQSAR_max3 | 0.90 | 0.00 | 0.49 | 0.49 | 159 | 136 | 0.86 |
| pQSAR_fp_yscram | 0.12 | 0.00 | 0.00 | 0.01 | 159 | 0 | 0.00 |
| pQSAR_0.05_yscram | 0.12 | 0.00 | 0.00 | 0.01 | 159 | 0 | 0.00 |
| pQSAR_0.2_yscram | 0.12 | 0.00 | 0.00 | 0.01 | 159 | 0 | 0.00 |
| pQSAR_max3_yscram | 0.12 | 0.00 | 0.00 | 0.01 | 159 | 0 | 0.00 |
| pQSAR_ensemble | 0.89 | 0.00 | 0.50 | 0.48 | 159 | 133 | 0.84 |
| pQSAR_ensemble_yscram | 0.12 | 0.00 | 0.00 | 0.01 | 159 | 0 | 0.00 |
| metaNN | 0.88 | 0.00 | 0.49 | 0.49 | 159 | 137 | 0.86 |
| metaNN_yscram | 0.05 | 0.00 | 0.00 | 0.01 | 159 | 0 | 0.00 |
| max4 | 0.90 | 0.00 | 0.53 | 0.51 | 159 | 140 | 0.88 |
| max4_yscram | 0.12 | 0.00 | 0.01 | 0.01 | 159 | 0 | 0.00 |
| ensemble_all | 0.91 | 0.00 | 0.54 | 0.52 | 159 | 139 | 0.87 |
| ensemble_all_yscram | 0.07 | 0.00 | 0.00 | 0.01 | 159 | 0 | 0.00 |

Here we also created an ensemble average of these 3 models, to adapt the benefit from multiple models, without the complication of multiple hypotheses. The simple equal weight averaged ensemble of these 3 models provided a marginally improved median R^2^_ext_ of 0.34 vs. 0.33 with the fixed correlation threshold 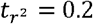. This ensemble did not change the median R^2^_ext_ (0.02) on the y-scrambled data set. In the following discussion and comparison with metaNN predictions, we use the predictions of this ensemble of the 3 pQSAR models as the “pQSAR result”.

For the same 4266 assays, the median R^2^_ext_ for metaNN is 0.33, almost exactly the same as obtained from the pQSAR ensemble (0.34). Similarly, metaNN had the same R^2^_ext_ in the y-scrambled data set of 0.02. The rank order plots in Figure 3 display R^2^_ext_ rank-order curves for metaNN and pQSAR, as well as the y-scrambled results. The overlay in the plots of metaNN and ensemble pQSAR demonstrates their comparable performance.

**Figure 3.**
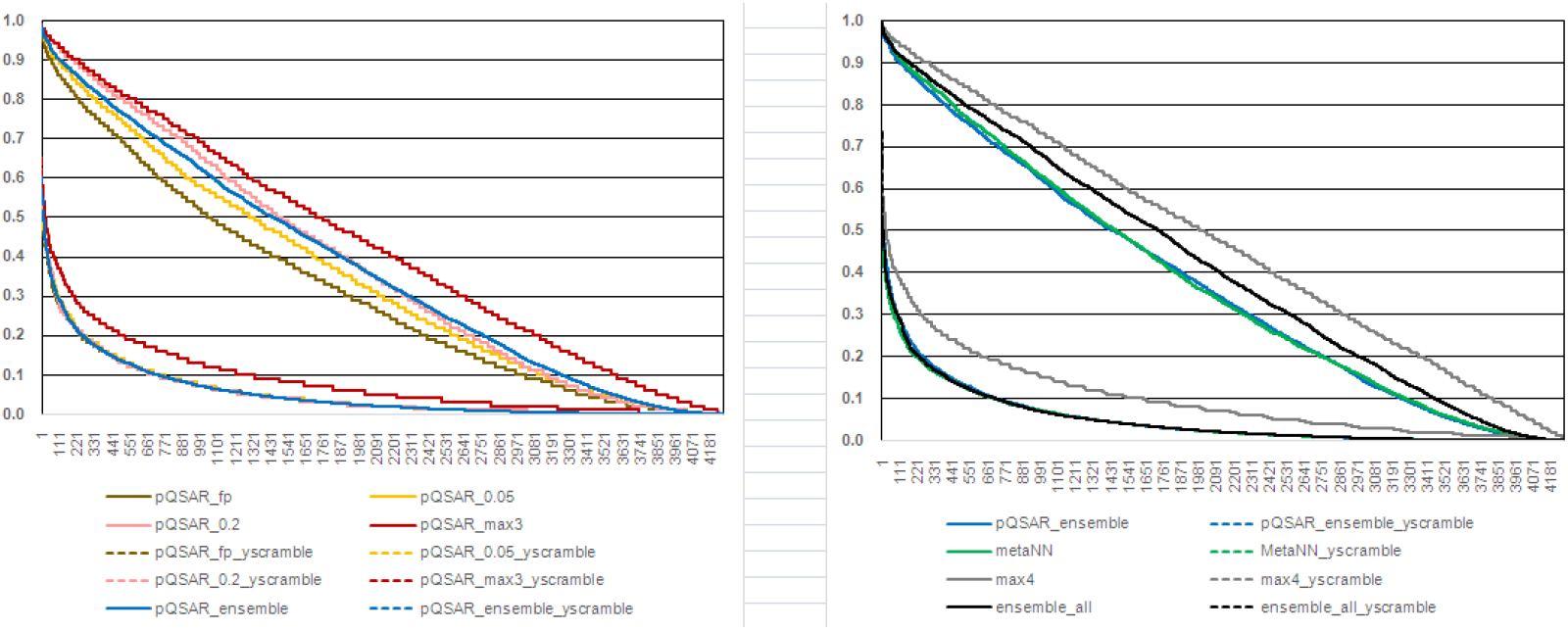
Left: pQSAR result; right: metaNN compared to ensemble pQSAR, and an ensemble of both methods.

We divided the 4266 assays into sub-groups to compare pQSAR and metaNN based on 4 criteria: protein families, assay size, assay format and activity distribution. Overall, the performance was comparable for different sub-groups. While each algorithm performed better for some types of assays, the difference was not large. As shown in Figure 4, each assay was assigned to 1 of 3 bins: equal for R^2^_ext_ difference between metaNN and pQSAR ensemble between -0.05 and 0.05; metaNN if the difference is > 0.05 and pQSAR if < -0.05. For many cases, the 3 bins have very similar numbers of assays. MetaNN performs slightly better for GPCR and cellular assays; ensemble pQSAR trains slightly stronger models for kinases. Both methods show a clear bimodal distribution with mean pIC_50_ in Figure 4d. However, pQSAR ensemble works better for assays with lower mean pIC_50_ around 5.0, whereas metaNN performs slightly better for assays with more potent compounds, peaking with mean pIC_50_ at 7.0. The dynamic range of pIC_50_ values as characterized by pIC_50_ standard deviation, and chemical diversity, as characterized by Bemis-Murcko scaffold count and cpd/scaffold count, were two additional factors used in comparison, but no substantial differences were observed.

**Figure 4.**
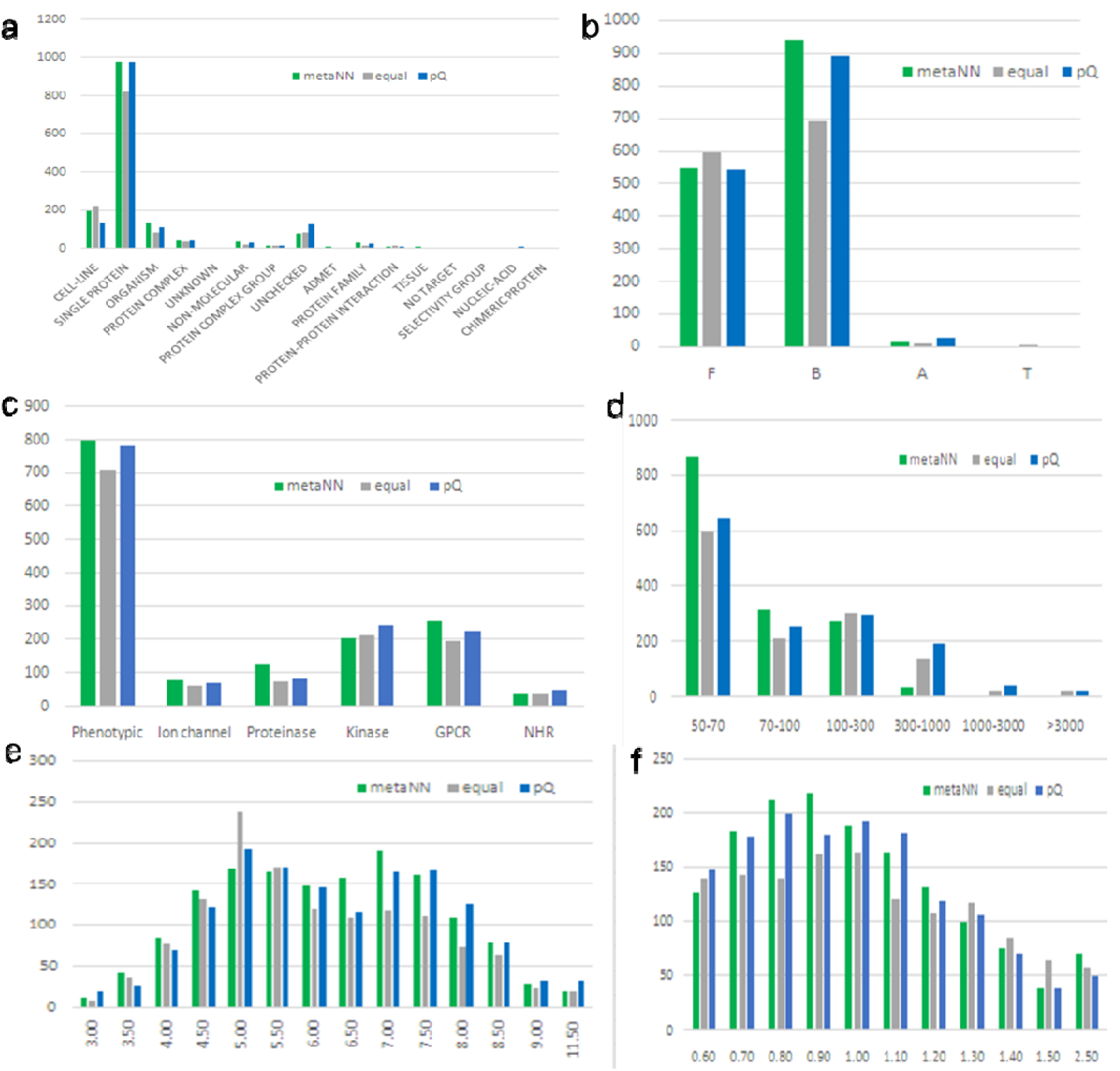
Prediction performance compared for metaNN and ensemble profile-QSAR, plotted for different sub-groups of assays. We note the R2ext difference between metaNN and pQSAR ensemble between -0.05 and 0.05 as equal and shown in the figures as gray bars; metaNN if the difference is > 0.05 and shown in the figure as green bars; and pQSAR if < -0.05 and shown in the figure as blue bars. a. Assay format and assay type from as extracted from ChEMBL database; b, A is the shortcut for ADMET assays, B for binding assays, F for functional assays, and T is for toxicity assays, c. Protein family: slightly different for kinase, protease and GPCR, d. Compound count: metaNN shows better performance for smaller assays, e. Mean pIC50 distribution, f. pIC50 standard deviation within each assay

One interesting observation is from the bins based on assay size. MetaNN gave overall higher R^2^_ext_ for smaller assays, especially those with no more than 100 compounds, while pQSAR ensemble performed better for the assays tested on 100-1000 compounds. This aligns with the expected strengths of metaNN and pQSAR. The high generalization capacity of the learned meta-initialization by metaNN enables the neural network to be quickly adapted to a target assay in only a few (i.e., 5 in our experiments) gradient steps with even a very limited number of training compounds. Profile-QSAR relies on more training compounds to build the PLS model, especially when the number of random forest models is high.

## 4. Discussion

We will first discuss appropriate measures of model “usefulness” for real-life drug discovery; then we will discuss the combination of different algorithms, the improved predictive ability from the combination, and whether there is a practical upper limit to prediction accuracy.

### 1). Realistic test set splitting and corresponding evaluating factors

Many machine-learning applications assume the training set is a fair sampling of the prediction set. However, drug-discovery teams are most interested in predictions for novel chemical matter very different from the previousl studied compounds used to train the models. A special procedure was therefore designed to split the whole data set into training and test sets specifically designed to mirror such “virtual screening”. Described in an earlier publication on pQSAR (*16*), the compounds from each assay are clustered based on their Morgan 2 chemical fingerprints. Models are trained on the 75% of compounds from the largest clusters and tested on the remaining small clusters and singletons. The test sets from this procedure are much less like the training set than the widely used random test sets. They were shown to mimic the novelty of compounds selected for testing in 18 real virtual screening projects, and thus are called the “realistic test sets”. When the performance of the models trained on this training set are evaluated against this test set, R^2^_ext_ should realistically indicate model performance in actual drug-discovery applications. In contrast, using random test sets dramatically overestimates prediction performance in real applications, which badly misleads project teams.

In real practice, training and test sets are subsequently combined, and a final model is trained on all the available bioactivities, which is then used to predict activity of new compounds with no experimental data. Still, assessing performance on the realistic test set is not just for comparing modeling methodologies. Project teams need these results to assess the expected accuracy of the models in practice--to know whether and how each model can best be employed in their projects. E.g., with a poorer model, screen a large number of predicted active compounds at a single concentration, followed by dose-response refinement on the experimentally confirmed hits, vs. with a better model, to save time and costs by immediately screening a smaller number of predicted actives with the dose-response assay. Similarly, consider model-quality in deciding which predicted bioactivities to follow up in off-target or mechanism-of-action calculations, to balance the probability of success with the costs of pursuing false-positives from virtually screening a large panel of models.

### 2). MetaNN and pQSAR can be profitably combined

As shown in the Results section, sometimes pQSAR and sometimes metaNN performs better for different assay sub-groups. We suggested that pQSAR can fish out complex combinations of relevant assays through the 2^nd^ level PLS modeling, but suffers from the curse of dimensionality, especially for smaller assays. MetaNN benefits more if the target assay aligns significantly with the consensus parameters. Since there is a theoretical reason for expecting one or the other to give better predictions for different assay types, we hypothesized that even better predictions might be achieved by a consensus of these very different algorithms. We tested two strategies for combining the predictions from 4 models (pQ_0.0, pQ_0.05, pQ_0.2 and metaNN): either by averaging the predictions, named “ensemble_all”, or by selecting the single model with the highest R^2^_ext_ on the realistic test set, named “max4”. Using the test set to choose among 4 models again raises the multiple-hypothesis problem. Note that the size of the problem increases with the number of hypotheses. An extensive optimization, such as a grid search over many parameters, would lead to extensive chance correlations. A simple selection among 4 models might produce few. There are numerous theoretical approaches and discussions around how to adjust parametric statistical measures for multiple hypotheses, but little consensus on the best practice. An alternative is to numerically measure the false discovery rate by repeating the calculation with scrambled pIC_50_s, as was done in the earlier pQSAR publication.

Both consensus approaches improved correlation with the test set more using real data than with Y-scrambled data, demonstrating the benefit of combining the methods. Median R^2^_ext_ using Max4 increased from between 0.26 and 0.33 for the 4 individual models, to 0.47 for selecting the best of the 4 for each assay, an improvement of from 0.14 to 0.21. Y-scrambled R^2^ _ext_ also increased, but only from 0.02 for the individual models to 0.07 for Max4. Similarly, the number of additional models with R^2^_ext_>0.3 found by max4 increased by 611 over the best single model (pQSAR using threshold 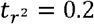), whereas the number of additional Y-scrambled Max4 models with R^2^_ext_>0.3 only increased by 159, for a net gain of 452 “successful” models. These improvements over chance support the conjecture that pQSAR and metaNN, although overall similar in individual performance, can be profitably combined.

For the “ensemble_all” consensus of averaging the 4 models, where no test set-based choices are made, the Y-scrambled median R^2^_ext_ remained at 0.02 as expected. With the real data, median R^2^_ext_ increased by 0.06 from 0.33 for the best single model to 0.39 for the ensemble average of all 4. Ensemble_all built 256 more models with R^2^_ext_>0.3 than the best single model, also with virtually no change in Y-scrambled results. Thus, Max4 explained additional 3% of the variance and built 196 additional successful models vs. ensemble_all. This improvement is modest but significant, since it summarizes results of over 4 thousand models. An additional advantage of Max4 is that predictions on large compound sets are faster, since ensemble_all requires 4 models per assay, and always includes pQSAR_0.0, by far the slowest for prediction.

In summary, both consensus approaches were improvements over a single model. The choice between ensemble_all and Max4 comes down to speed, accuracy and risk tolerance. The end-user must ultimately decide if the improved speed, larger number of successful models and higher average prediction accuracy of Max4 over the ensemble average is worth the slightly higher chance of being misled by a chance correlation.

### 3). Is there an upper limit of prediction accuracy?

MetaNN and pQSAR individually show very similar prediction performance calculated by R^2^ _ext_, for the 4266 data sets. We have also compared the models trained on the set of 4266 diverse assays to those from the much smaller and more homogeneous set of 159 kinase assays. We again calculated the 3 pQSAR models using thresholds of 0.0, 0.05 and 0.2, pQSAR_max3, pQSAR ensemble, metaNN, Max4, and ensemble_all consensus models, along with their y-scrambled controls. Table 3 summarizes the performance statistics. As expected, with much more homogeneous data, the transfer of learning, and thus the models for these kinase assays, are overall much stronger. The median R^2^_ext_ values are all above 0.45. The results also maintain the same trends as the 4266 assay results discussed above: pQSAR ensemble is comparable to metaNN, both with median R^2^_ext_ of 0.49-0.50, max4 and ensemble_all of metaNN and pQSAR further improve the median by 0.04, while the Y-scrambled result was almost unchanged. Although the trends are the same, the marginal improvement was significantly smaller. One underlying reason might be that the chemical diversity involved in the data set is much smaller. Another possible reason is that all targets share similar ATP binding sites, so all assays are correlated and contribute signal for pQSAR, and all are near the consensus model for metaNN. For these well-studied targets, both methods produced strong models for a much larger portion of the assays, with up to 88% of assays producing “successful” models with R^2^_ext_ >0.3. This raises the question: would there be an upper bound of this prediction accuracy, and if so, how high is it?

## 5. Conclusion

We presented a novel transfer-learning algorithm, metaNN, and reported its prediction performance on two public-domain bioactivity data sets. The performance was found to be surprisingly comparable to and even marginally better than the earlier pQSAR algorithm. Besides, it enjoys greater storage efficiency; a consensus DNN model is stored only instead of all previous random forest models required by pQSAR. We demonstrated that combining these two very different algorithms further improves the predictions: a 27% increase of median R^2^_ext_, and successful models for another 10% of the >4000 assays. We anticipate these newly developed algorithms will have wide applications in drug discovery. Future directions include how incorporating assay or target meta-data might further improve the models, and might also help to elucidate the relationships among experimental assays, e.g. mechanisms of action for the phenotypica assays.

There are many other multitask algorithms applicable to ligand-based bioactivity prediction: alchemite, multitask fully-connected deep neural networks, proteochemometrics and content-based filtering are a few. The authors are working with collaborators to analyze results from these methods for a more extensive analysis.

## Acknowledgements

Dr. Li Tian acknowledges “Laboratory for Synthetic Chemistry and Chemical Biology” under the Health@InnoHK Program launched by Innovation and Technology Commission, The Government of Hong Kong Special Administrative Region of the People’s Republic of China, for the funding support of this research.

